# Deciphering Molecular Cascades in a Novel Acclimatization Strategy for Rapid Ascent to High Altitude

**DOI:** 10.1101/145342

**Authors:** Subhojit Paul, Anamika Gangwar, Kalpana Bhargava, Yasmin Ahmad

**Affiliations:** Defence Institute of Physiology & Allied Sciences, Defence R&D Organization, Timarpur, New Delhi-110054

**Keywords:** Proteomics/Network, biology/cytoskeleton/redox, homeostasis/rapid, acclimatization

## Abstract

The repercussions of hypobaric hypoxia are dependent upon two factors-time and intensity of exposure. The effects of intensity i.e. variation of altitude are yet unknown although it is a significant factor in terms of acclimatization protocols. In this study we present the effects of acute (24 h) exposure to high (10,000 ft), very high (15,000 ft) and extreme altitude (25,000 ft) zones on lung and plasma using semi-quantitative redox specific transcripts and quantitative proteo-bioinformatics workflow in conjunction with redox stress assays. Our findings indicate that very high altitude exposure elicits systemic redox homeostatic processes due to failure of lung redox homeostasis without causing mortality. We also document a rapid acclimatization protocol causing a shift from 0 to 100% survival at 25,000 ft in male SD rats upon rapid induction. Finally we posit the various processes involved and the plasma proteins that can be used to ascertain the acclimatization status of an individual.

## INTRODUCTION

Around 466 million humans reside permanently above 3000 msl in Ethiopia (East Africa), Tibet (Asia), Andes (South America) and elsewhere around the globe[1, 2]. This is above and beyond the annual influx of tourists, athletes and adventurers. Hence, acclimatization or the lack of it, even when not considering the related high-altitude maladies, is a significant socio-economic process in high-altitude areas. Altitude has been categorized arbitrarily into three zones: High altitude (5,000–11,500 ft); Very high altitude (11,500–18,000 ft) and Extreme altitude (>18,000 ft)[3]

The majority of altitude sickness cases have been reported at altitudes >8,000 ft[4–6]. The altitudes ≥25,000 ft are considered as “Death zone” as no human body can acclimatize at that altitude. It serves as the upper limit of acclimatization to altitude. Although low temperature, low humidity and inhospitable terrain constitute a part of the collective complications at high-altitude, the basic cause attributed to high altitude maladies is hypobaric hypoxia[7–9].

Hypobaric hypoxia exposure is two-pronged-time of exposure (temporal variation, i.e. acute vs chronic) and intensity of exposure (altitude variation, i.e. high vs very high vs extreme). As per our knowledge, this is the first attempt to understand the effects of altitude variation on the rat lung and plasma proteome in conjunction with biochemical redox stress parameters. Most of the previous studies have focused on the temporal effects of a sample set at a particular altitude regarding acclimatization with most authors discouraging a high rate of ascent[10–12]. Some articles have also discussed the ascent of similar climbers in similar mountain ranges but at different ascent rates with suggestions for identification of susceptible and prophylaxis[13–17]. But the process of acclimatization remains untouched in all the articles cited above let alone any suggestions regarding rapid acclimatization. The process of acclimatization as followed by most mountaineers consists of slow graded ascent. But even during and after acclimatization, there are cases of AMS, HACE and HAPE reported[18–21]. The diagnosis of these maladies is mostly dependent on qualitative physiological parameters. The Lake Louise Score for AMS, for example, provides a score based on symptoms such as nausea, quality of sleep and headache, which to a certain extent are subjective parameters based on individual assessment[22]. We therefore confront two long-standing issues related to high-altitude acclimatization. The first is of rapid induction to high altitude. The second, consequently, is the ability to decipher the acclimatization status objectively using a panel of plasma proteins. Hence, we ask if the intensity of hypobaric hypoxia, i.e. altitude variation for acute period provides an alternative method for rapidly acclimatizing to a higher altitude. We have checked the effects of acute (24 h) exposure to simulated altitudes of 10,000 ft (high altitude zone), 15,000 ft (very high altitude zone) and 25,000 ft (extreme altitude zone) on SD rats at the proteome and oxidative stress specific transcripts levels. Our previous work stating the feasibility and statistical robustness of Sult1A1 as a marker for hypobaric hypoxia stress (and HAPE) and 24 h being the unique time-point showing maximal perturbations within the proteome[23] are also being re-invoked here in the context of altitude variation. Consequently, based on molecular cascades, can there be preventive diagnosis for establishing objectively if an individual is acclimatizing or not based on molecular events much before life-threatening physiological consequences of hypobaric hypoxia occur. It will be a more objective and quantitative method for determining individuals’ acclimatization status in association with the already established subjective and qualitative tests (e.g.-Lake Louise Criteria) further augmenting them.

In present study we report that acute exposure to very high altitude (15,000 ft) causes maximal tolerance to hypobaric hypoxia at the proteome and biochemical levels thereby causing a 100% shift from mortality to survival during further exposure to extreme altitude of 25,000 ft. We also observe lung redox homeostasis is completely voided at very high altitudes leading to activation of systemic redox homeostasis. But upon revival of lung redox homeostasis in 25K (A) group due to the acclimatization strategy we observe subdued systemic redox homeostasis. Herein, we also state plasma proteins that can indicate an organism’s redox status at high, very high and extreme altitude thereby objectively assess its state of acclimatization.

## RESULTS

### Survivability and Mortality upon induction to extreme altitude (25,000 ft): Pre-exposure to15,000 ft followed by normobaric normoxia exposure

Five groups of SD rats were exposed to Normobaric normoxia/Baseline (BL: 744 ft); 10,000 ft (10K, high altitude zone); 15,000 ft (15K, very high altitude zone) and 25,000 ft (25K, extreme altitude zone) as shown in Fig. 1. The ascent rate was kept very rapid (589 m/min). Two groups were exposed to 25,000 ft- one group, 25K (D) was exposed directly to 25,000 ft without any acclimatization while the other group, 25K (A) went through a very short acclimatization (10 h at 15,000 ft followed by 1 h of normobaric normoxia before ascent to 25,000 ft). It was observed that till 15,000 ft there was no mortality while all rats given direct exposure to 25,000 ft, i.e. 25K (D) group perished (Fig.1). But upon pre-exposure to 15,000 ft (simulated) for 10 h with subsequent exposure to normobaric normoxia for 1 h prior to exposure of 25,000 ft (simulated), there was a complete reversal of mortality to 100% survivability, as observed in the 25K (A) group. This phenomenon prompted us to delve into the molecular events causing reversal of mortality to survivability.

**Figure 1:**
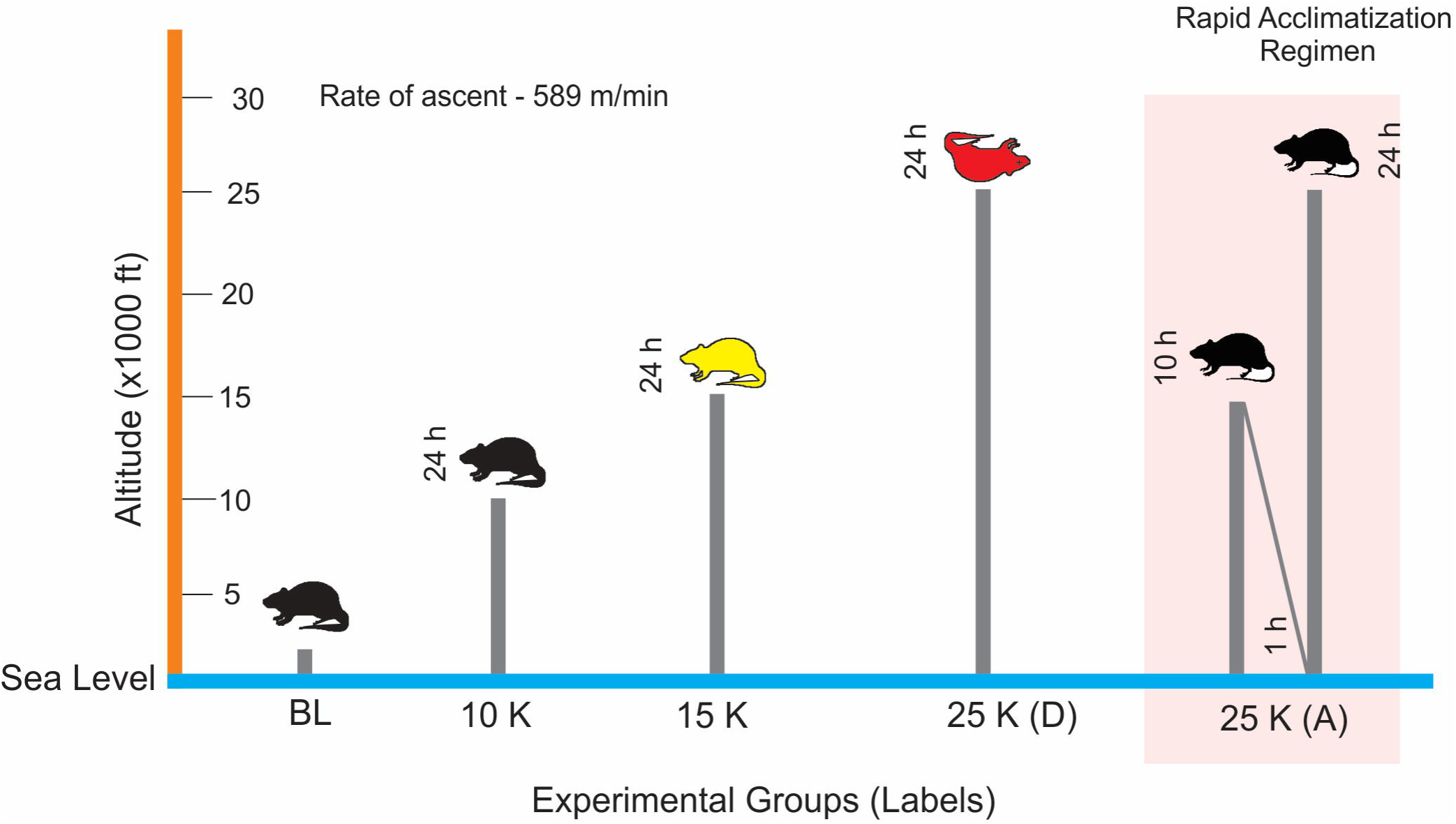
Study Design. Male SD rats (10 week old; 200–230 g) were exposed to Normobaric normoxia (BL; mmHg; n=9), 10,000 ft (10K; mmHg; n=9), 15,000 ft (15K; mmHg; n=9), 25,000 ft (25K; mmHg; n=18). Exposure to 25,000 ft was provided to two separate groups. One group (25K (D); n=9) was exposed directly to 25,000 ft altitude while the other group (25K (A); n=9) was given an acclimatization for 10 h at 15,000 ft and 1 h at normobaric normoxia. All altitude exposures were simulated in hypobaric hypoxia chamber with the rate of ascent being 589 m/min and temperature (25˚C) and humidity (50%). Zero animals survived in 25K (D) group while all animals (9) survived in 25K (A) group. Upon biochemical and omics-based investigation on lung tissue and plasma, we observed 15K group (yellow) had maximum perturbations in redox and energy homeostasis with failing housekeeping functions in lung tissue while all other groups (black) had minimal/reduced molecular perturbations. 25K (D) group (red) could not be probed as all animals in this group died on exposure. The experiment was repeated thrice for statistical significance with each group having three rats per experimental replicate.

### Rapid induction causes activation of acute phase signaling and impairment of redox stress management in lung tissue while activating the systemic redox homeostatic mechanisms in 15K group: Benefits in 25K (A) group

Since the lung tissue is the primary interface between atmosphere and organism, we began our investigation with the lung proteome (iTRAQ labeled; LC-MS/MS). A total of 117 (10K group), 92 (15K group) and 123 (25K (A) group) proteins were identified (Fig.2a). But in 15K group there is unanimous down-regulation with the number of proteins identified being reduced noticeably. 25K (A) group to the contrary displays the maximum number of up-regulated proteins and minimum number of down-regulated ones. Common proteins between all the three groups from lung tissue was just one while 13 proteins were common between 10K and 15K groups, 8 proteins were common between 10K and 25K (A) and 23 proteins were common between 15K and 25K (A) groups (Fig.2b). Since we found a lot of proteins in the lung proteome to be correlated to oxidative stress, a PCR array analysis of lung tissue for oxidative stress markers (*Supplementary dataset S1*) was performed. The clustergram of which revealed that 25K (A) group was closer in trend to 10K and normoxic groups while opposite to 15K group (Fig. 2c).

**Figure 2:**
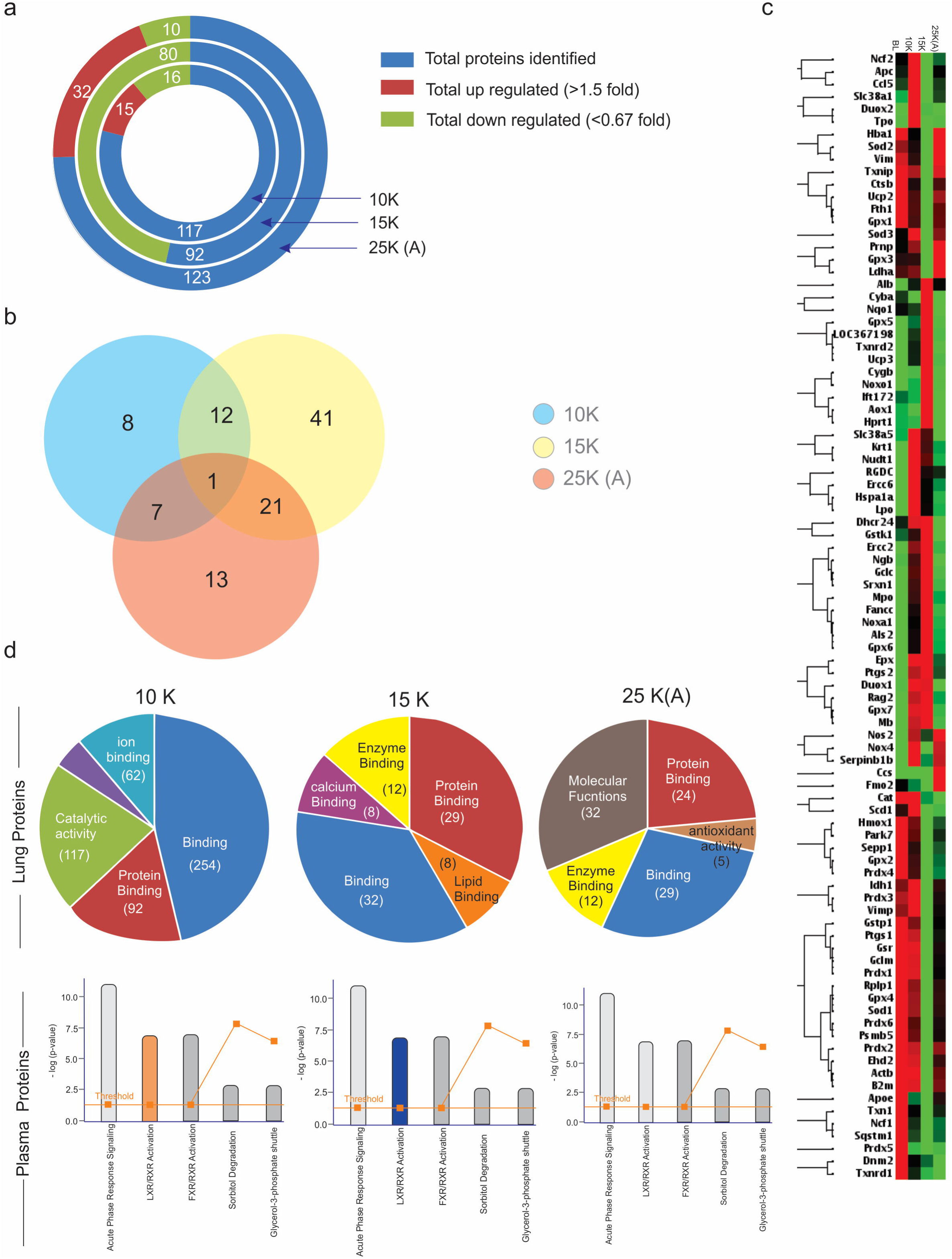
Overview of Proteome and redox-specific transcripts alongwith affected pathways. a. The total number of proteins (blue) identified in 10K, 15K and 25K (A) groups alongwith number of up-regulated (maroon) and down-regulated (green) proteins during LC-MS/MS (iTRAQ labeled) analysis of lung tissue. 15K group has the least number of proteins identified (92) with the maximum number of down-regulated proteins (80). 25K (A) group had the highest number of proteins identified (123) among which 32 were up-regulated and 10 down-regulated. 10K had 117 proteins identified with 15 up-regulated and 16 down-regulated proteins. Fold change value greater than 1.5 fold was considered up-regulation while fold change value lesser than 0.67 was considered as down-regulation.
b. Venn diagram showing overlapping up- and down-regulated proteins among the proteins in 10K (blue), 15K (yellow) and 25K (A) (pink) groups. Fold change values of 1.5 or more and 0.66 or less were the criteria for choosing up-regulated and down-regulated proteins respectively. Venn diagram was created using Venny 2.1 (*Oliveros, J.C. (2007–2015) Venny. An interactive tool for comparing lists with Venn’s diagrams.* http://bioinfogp.cnb.csic.es/tools/venny/index.html). Only a single protein was common between 10K, 15K and 25K (A) groups. The maximum number of overlapping proteins was between 15K and 25K (A) groups (22). The minimum number of overlapping proteins was between 10K and 25K (A) (08).
c. Clustergram with maximum join type hierarchical clustering comparing the redox-stress specific transcripts’ up-regulation (red) and down-regulation (green) among BL (baseline-normoxic control), 10K, 15K and 25K (A) groups. PCR-Array specific to redox-stress specific transcripts was used to generate ΔΔC_T_ values (RT-PCR) for all four groups (BL, 10K, 15K & 25K (A)) which were further analyzed using RT^2^ Profiler PCR Array Data Analysis Version 3.5 from Qiagen. 15K and 25K (A) group shown anti-trends while 25K (A) trends for transcripts match those observed in 10K and BL.
d. Gene ontology analysis of lung proteome revealed that Binding and Protein binding processes dominate across all three groups while 25K (A) group has molecular functions as the top-most molecular function. Pathway analysis (IPA) of plasma proteins reveals that LXR/RXR Activation is perturbed in 10K (up-regulated; saffron; z-score>2) and 15K (down-regulated; blue; z-score<2) groups while its normalized in 25K (A) (grey; z-score either zero or nonsignificant) group. Acute phase signaling remains the pathway with the maximum number of proteins across all three groups. A co-relation is indicated between the molecular functions in lung proteome and the perturbed pathway in plasma proteome.

GO analysis of the lung proteome revealed binding and protein binding processes to be common and dominant across all three groups except in case of 25K (A) group where molecular functions dominate (Fig.2d). A mix of high-throughput and low-throughput modules was used for the analysis of lung and plasma proteomes, using LC-MS/MS (iTRAQ labeled) (*Supplementary dataset S2*) and MALDI-TOF/TOF (*Supplementary dataset S3*), respectively. While LC-MS/MS provided a huge list of proteins with quantification to understand the underlying causal events in lung, the whittling down of this list was achieved in plasma proteome by MALDI-TOF/TOF. Upon pathway analysis (using IPA) of the plasma proteins, we observed Acute phase response signaling (APRS) and LXR/RXR Activation to be the two most significant pathways (Fig.2d). LXR/RXR Activation was up-regulated in 10K (z-score>2); down-regulated in 15K (z-score<-2) while it showed no perturbations in 25K (A) group (z-score=0). Overall, all these findings indicate a strong impetus towards a systemic stress at 15K due to redox imbalance and associated processes.

In Table 1, down-regulation (fold change≤0.6) of CRP, APO H, Fibrinogen alpha and gamma chains, VDBP etc in 15K group plasma but a significant increase (fold change value≥1.5) of glutathione peroxidase 3 and hemopexin was observed. Serum albumin, a negative acute phase protein, showed decreased levels (0.63 fold) in 15K group before recovering (1.12 fold) in 25K (A) group.

**TABLE 1:** 2D-MALDI-TOF/TOF identified proteins with Uniprot ID and normalized fold change values across each group.

| S.No. | UniprotID | Protein ID | Normoxia | 10K | 15K | 25K(A) |
| --- | --- | --- | --- | --- | --- | --- |
| 1 | P02770 | Serum albumin | 1 | 0.69564562 | 0.632728273 | 1.123398116 |
| 2 | J3QLI0 | Beta-2-glycoprotein 1 APO H | 1 | 0.599191064 | 0.442797632 | 0.98672265 |
| 3 | P20059 | Hemopexin | 1 | 1.078078586 | 1.702732484 | 1.580064789 |
| 4 | P04276 | Vitamin D-binding protein | 1 | 1.099381524 | 0.614069314 | 0.951075257 |
| 5 | P02680 | Fibrinogen gamma chain | 1 | 0.506130817 | 0.343636037 | 1.268732041 |
| 6 | P48199 | C-reactive protein | 1 | 0.960973541 | 0.178058842 | 0.677460411 |
| 7 | P23764 | Glutathione peroxidase 3 | 1 | 1.451303161 | 8.596477529 | 1.390318158 |
| 8 | P02767 | Transthyretin | 1 | 0.864864807 | 0.598442174 | 0.871346305 |

There were common links in terms of pathways (acute phase and LXR/RXR signaling) and directly in terms of molecules (hemopexin, serum albumin, glutathione peroxidase 3) between the lung proteome and plasma proteome across different proteomics platforms suggesting a certain “unity in diversity” for the platforms and protein sets. As oxidative stress is the principal event of hypobaric hypoxia exposure, we analyzed the ROS and MDA levels in both lung and plasma (Fig.3a & 3b). We observed that although both lung and plasma had the same trend in ROS generation, MDA levels were quite different. MDA levels (indicator of oxidative damage) were highest in 15K group lungs (□5 fold) while plasma samples had the highest MDA levels in 10K group (□1.5 fold). Both lung and plasma had lower MDA levels in 25K (A) group. The results suggest that although the ROS generation continued with hypobaric hypoxia exposure, it was neutralized systemically in 15K plasma but caused damage resulting in secondary products like MDA in 15K lungs. IPA analysis based extrication of APRS revealed that levels of proteins with anti-oxidant roles as well as roles in inflammatory axis were perturbed (Fig.3 c). MCP-1(monocyte chemotactic protein-1) is a chemoattractant for causing the extravasation of circulating monocytes and macrophages out of the blood vessels’ endothelium. It subsequently causes inflammation at the site of extravasation[24, 25]. Thus we measured MCP-1 in plasma to assess systemic inflammatory stimulus as per altitude variation (Fig.3d). We observed its levels skyrocket in 10K (0.6 ng/ml) and further in 15K (0.91 ng/ml) group while 25K (A) had negligible levels (0.18 ng/ml). This indicates remarkable reduction of monocyte infiltration, the causal event of inflammation[26, 27]. In line with the quest for putative markers we analyzed Sult1a1 levels also (Fig.3e) as we had observed it had statistically significant discriminatory powers earlier[23]. In the context of altitude variation also, Sult1a1 proved to be a good indicator of acclimatization as indicated by its similar levels in plasma of normoxic controls and 25K (A) group (□1500 µmol/l).

**Figure 3.**
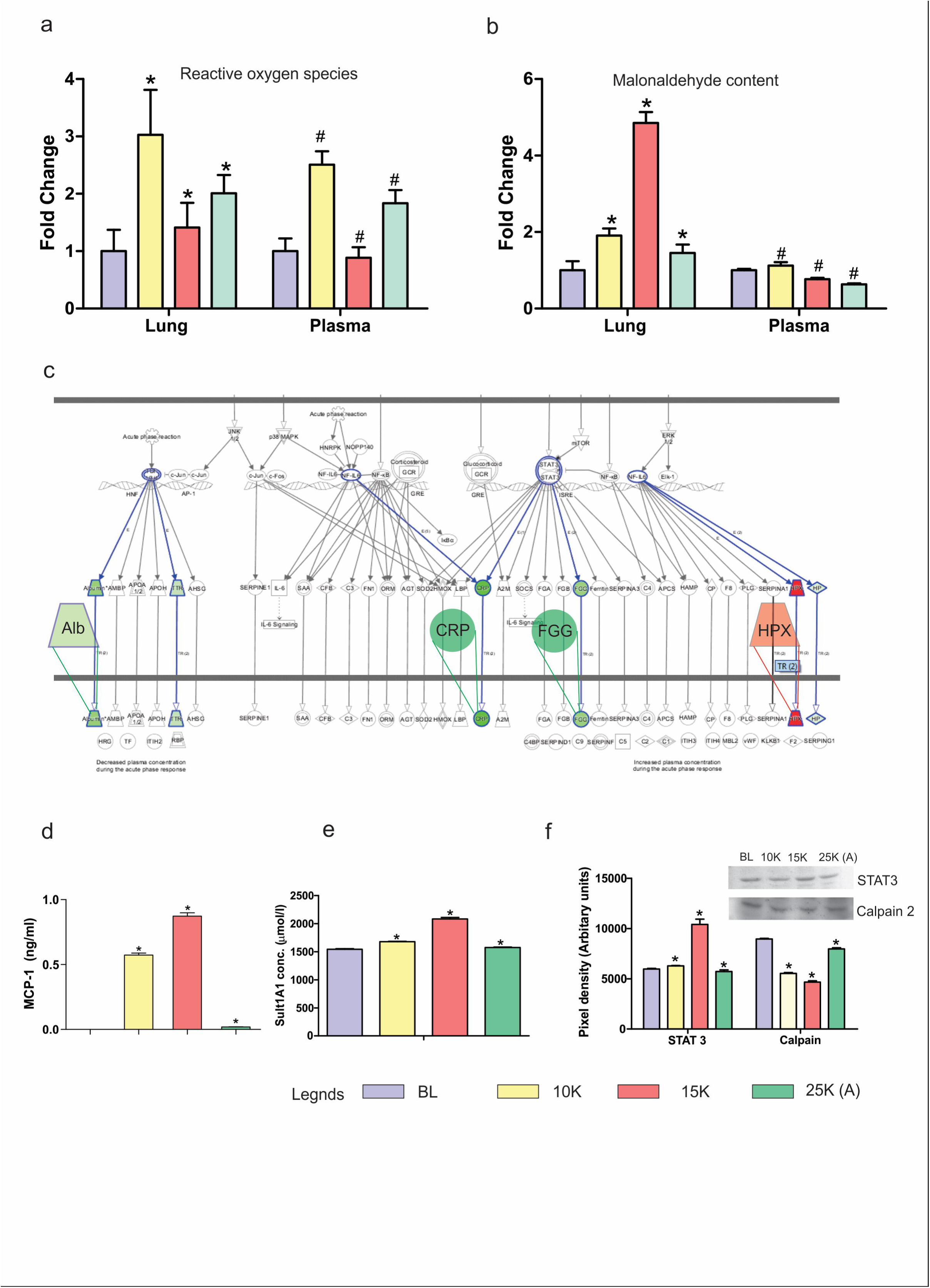
Redox processes and Acute phase signaling. a. ROS levels were measured in both lung and plasma using DCFH-DA assay. The assay was performed in triplicates with three readings per group per experiment. ROS levels in both lung (fold change 3) and plasma (fold change 2.5) increase in 10K, decrease in 15K (Lung: fold change 1.5; Plasma: fold change 0.8) and finally increase marginally in 25K (A) (Lung: fold change 2; Plasma: fold change 1.8). Results are presented as Mean±SEM. Mean was calculated from the three separate experimental replicates. * represents p-value<0.0001 in lung tissue while # represents p-value<0.0001 in plasma.
b. MDA (malondialdehyde) levels were measured in both lung and plasma using TBARS assay. In lung, MDA levels increase slightly in 10K group (fold change 2), become highest in 15K group (fold change 5) and decline significantly in 25K (A) group (fold change 1.6). In plasma, there is a slight increase in 10K group (fold change 1.3) before subsequently declining in 15K (fold change 0.7) and 25K (A) (fold change 0.5). Results are presented as Mean±SEM. Mean was calculated from the three separate experimental replicates. * represents p-value<0.0001 in lung tissue while # represents p-value<0.0001 in plasma.
c. Expanded view (representative of 15K group) of IPA mined Acute phase signaling pathway with overlaid up-regulated (red) and down-regulated (green) proteins identified in plasma proteome (Supplementary dataset S3).
d. Monocyte chemoattractant protein-1 (MCP-1) levels were measured in plasma. It is an indicator of monocyte activity and inflammatory stimuli. MCP-1 levels increased sharply in 10K (0.6 ng/ml) and 15K (0.8 ng/ml) before declining sharply in 25K (A) (0.1 ng/ml). Results are presented as Mean±SEM. Mean was calculated from three separate experimental replicates. * represents p-value<0.0001.
e. Sulfotransferase 1A1 (Sult 1A1) levels were measured in plasma. It is an indicator of hypoxic stress and found elevated in HAPE patients’ plasma[23]. ELISA results show it increases in 10K (1600 µmol/l) and 15K (2200 µmol/l) as compared to Baseline (BL; 1500 µmol/l) while levels in 25K (A) group are almost equal to BL. Results are presented as Mean±SEM. Mean was calculated from the three separate experimental replicates. * represents p-value<0.0001.
f. Representative immunoblots of STAT-3 and Calpain-2 in plasma with bar-graphs showing differential levels of both proteins. STAT-3 levels increase drastically in 15K group (10,000 AU) while other groups have similar levels (approx. 5,000–7,000 AU). Calpain-2 decreases in 10K (6,000 AU) and 15K (4,000 AU) group as compared to BL (9,000 AU) with significant recovery in 25K (A) group (8,000 AU). The anti-trends observed in STAT-3 and Calpain-2 levels across the four groups indicate a systemic transcriptional activation and down-regulation of proteolysis that reaches its peak in 15K group and is reverted to BL levels in 25K (A) groups. Results are presented as Mean±SEM of autoradiograms’ pixel intensities (Arbitrary units). Mean was calculated from three separate experimental replicates. * represents p-value<0.0001.

Finally, we also examined the levels of upstream APRS molecules (STAT-3) and effectors (calpain-2) in plasma samples (Fig.3 f). Immunoblotting revealed STAT-3 levels are highest in 15K group plasma (□ 10,000 AU) indicating a perturbed signaling cascade in 15K group while 25K (A) group (□ 5,000 AU) showed modest decline as compared to 15K group. In calpain-2 the levels are similar in normoxia and 25K (A) group (□9000 AU) indicating reduction in protein misfolding (indicator of oxidative stress) in 25K (A) group. It has roles in signal transduction, pulmonary vascular remodeling and cytoskeletal remodeling as well[28–30].

As shown in Figures 2 & 3, 15K group is the group where there is an offsetting of the redox balance in lung tissue and the activation of systemic redox stress management mechanisms to ensure survival of the organism. But unlike 25K (D) group, the 15K group had no mortality suggesting activation of the systemic mechanisms against redox stress without causing a mortal threat to the organism.

### Perturbed antioxidant reserves in 15K group were restored in 25K (A) group: Rapid induction is a systemically feasible process

Since the oxidative stress response occupied centre stage in lung proteome and transcriptome, we investigated Nrf2 levels in lung (Fig.4a). As was expected, Nrf2 (which mediates oxidative stress response and helps the cell survive) declined in 15K group (□ 11,000 AU) while 25K (A) group showed restoration to normoxic levels (□ 17,000 AU). A downstream effector of the Nrf2 mediated response, peroxiredoxin 6 (Prdx6) also showed abysmal decline in 15K group (□ 1,000 AU) before recovering in 25K (A) group (□11,000 AU) (Fig.4a). Furthermore plasma levels of GPx3 and TR2 were also investigated. GPx3 increased (□ 27,000 AU) when most other antioxidant enzymes were in decline while TR2 exhibited a decline in 15K group (□7,000 AU) and recovery in 25K (A) (□10,000 AU) coinciding with normalization of GPx3 levels (□11,000 AU) (Fig.4a). The direct hints in the lung and plasma proteomes regarding the related antioxidant reserves of glutathione (GPx3 in MALDI and LC-MS/MS) and thioredoxin (thioredoxin levels in LC-MS/MS) points to the important, almost essential roles these two had to play in maintaining systemic redox homeostasis. This is also evident in the fact that loss-of-function mutations of thioredoxin are lethal to the zygote[31–33]. LC-MS/MS data showed thioredoxin levels decrease in 15K dramatically (0.126 fold) and increase sharply in 25K (A) group (3.158 fold) in lung tissue. TR2 is an enzyme that keeps thioredoxin in reduced state[31]. We observed that its levels were mirroring Trx levels systemically.

**Figure 4.**
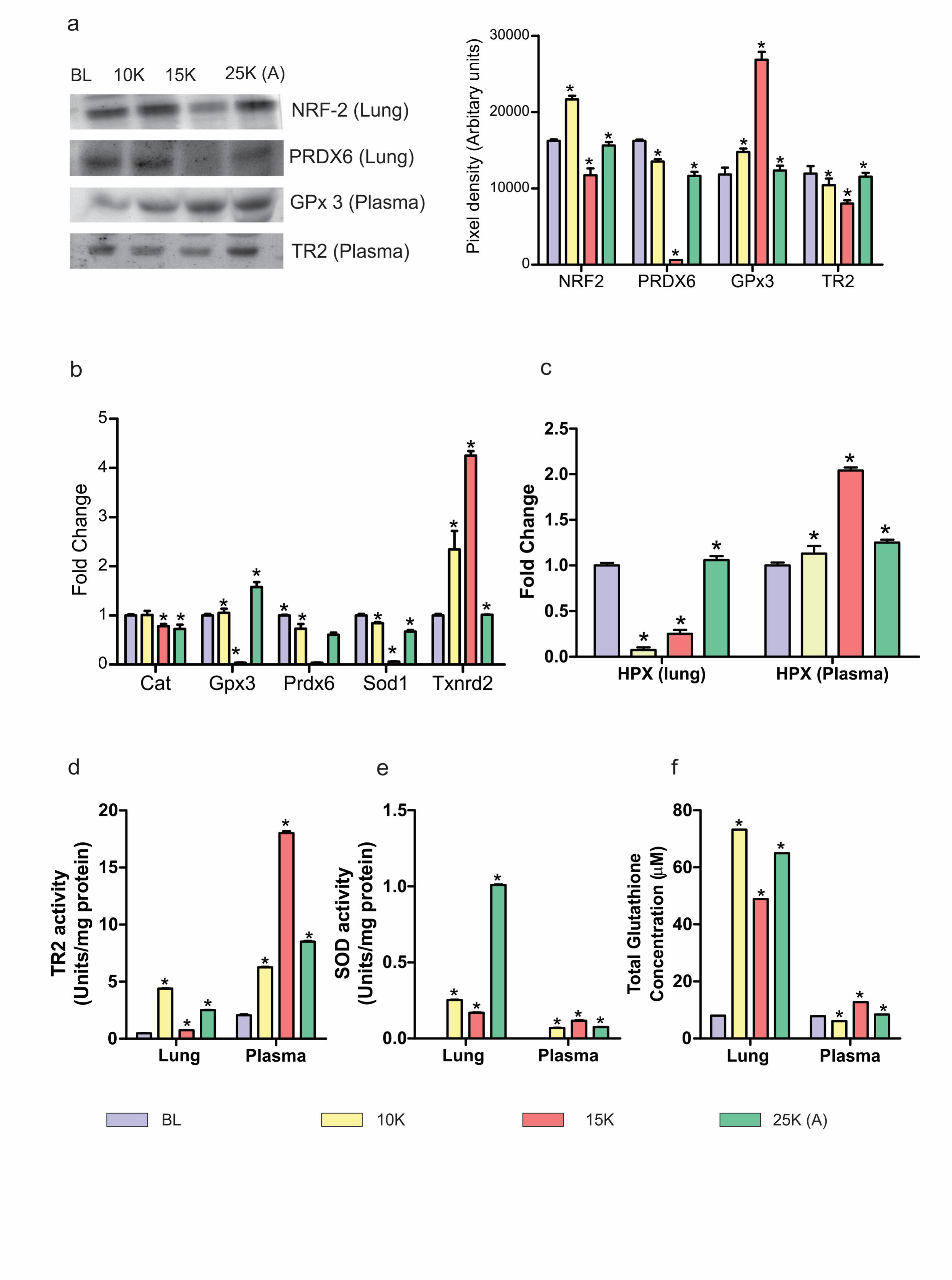
Nrf2 signaling in lung and its downstream systemic effects. a. Representative immunoblots of Nrf2 and PRDX6 in lung tissue and of GPX3 and TR2 in plasma with bar-graph depicting their levels in each group. Lung tissue: Nrf2 levels increase in 10K (22,000 AU), decrease in 15K (12,000 AU) and revert to BL levels (16,000 AU) in 25K (A) group. PRDX6 slightly decrease in 10K (14,000 AU), decline drastically in 15K (1000 AU) and recover sharply in 25K (A) (11,000 AU) as compared to BL (17,000 AU). Plasma: GPX3 levels increase slightly in 10K (14,000 AU), increase further (27,000 AU) and decline to close to normoxic levels (11,000 AU) in 25K (A) group. TR2 levels decline noticeably in 15K group (7,000 AU) and recover close to normoxic levels (11,000 AU) in 25K (A) group. Results are expressed as Mean±SEM of autoradiograms’ pixel intensities (Arbitrary units). Mean was calculated from three separate experimental replicates. * represents p-value<0.0001.
b. Fold change values of select redox-stress specific transcripts from PCR-Array performed using lung tissue. Across major antioxidants’ transcripts 15K group samples show declining levels, except in Txnrd2 (TR2) which shows increased fold change in 15K group.
c. ELISA was performed on lung tissues and plasma to assess Hemopexin levels. In lung tissue, hemopexin declines in 10K (0.1 fold change) and 15K groups (0.25 fold change) and rebounds to normoxic levels (1.0 fold change) in 25K (A) group. In plasma, there is significant increase in hemopexin levels in 15K (2.1 fold change) before levels decline in 25K (A) (1.3 fold change) Bar graph depicts results from each group as Mean±SEM. Mean was calculated from three separate experimental replicates. * represents p-value<0.0001.
d. Bar graph depicting thioredoxin reductase 2 (TR2) activity levels in each group across lung tissue and plasma. In 15K group, lung tissue show maximum decline in TR2 activity (1 unit/mg protein) while plasma has highest activity (19 units/mg protein). In 25K (A) group, TR2 activity resurges in lung (3 units/mg protein) but declines in plasma (8 units/mg protein). Results are depicted as Mean±SEM. Mean was calculated from three separate experimental replicates. * represents p-value<0.0001.
e. Bar graph depicting Superoxide dismutase (SOD) activity levels in each group across lung tissue and plasma. In 15K group, lung tissue witness a decline in SOD activity (0.2 units/mg protein) while plasma shows increased SOD activity (0.2 units/mg protein). In 25K (A) group, lung tissue has highest SOD activity (1.0 unit/mg protein) while plasma SOD activity declines (0.1 unit/mg protein). Results are depicted as Mean±SEM. Mean is calculated from three separate experimental replicates. * represents p-value<0.0001.
f. Bar graph depicting total glutathione (GSH) concentration in all groups across lung tissue and plasma. In lung tissue, GSH levels increase in 10K (75 µM), decrease in 15K (47 µM) and increase again in 25K (A) (65 µM). In plasma, GSH levels decrease slightly in 10K (5 µM), increase in 15K (13 µM) and fall back close to normoxic levels (7 µM) in 25K (A). Results are depicted as Mean±SEM. Mean was calculated from three separate experimental replicates. * represents p-value<0.0001.

The PCR array data for lung tissue also reinforces the fact that 15K group had the maximum down-regulation of anti-oxidant proteins (Fig.4b) as seen in GPx3, Prdx6 (fold change≤0.3) & Txnrd2 (TR2) (fold change □5). Here we found that there were opposite trends between Txnrd2 and others. This again points towards activation of systemic redox homeostatic mechanisms upon failure of lung redox homeostasis, possibly via Nrf2. To investigate this aspect, we investigated central antioxidants like Hpx (Fig.4 c), TR2 (Fig.4d), SOD (Fig.4e), GSH (Fig.4 f) in both lung and plasma. For all, we observed opposing trends in 15K and 25K (A) group in lung and plasma. In lung, if there were decreases in 15K group (Hpx=0.25 fold change, TR2=0.1 units/mg, SOD=0.2 units/mg, GSH=50 µM); 25K (A) group definitely shows rebound (Hpx=1 fold change, TR2=3 units/mg, SOD=1 unit/mg, GSH=65 µM). The same is also true for plasma in case of 15K and 25K (A) group. To be concise, plasma antioxidants normalize to base-line levels if the lung anti-oxidants recover (GSH, Hpx) but when the lung anti-oxidants can’t recover (SOD) systemic levels of that anti-oxidant remain high to supplement its function. Thus, the systemic redox homeostasis remains effective and circumvents mortality.

### Cytoskeletal re-arrangements & Perturbed housekeeping proteins: Massive down-regulation in 15K group but recovery in 25K (A) group

Nrf2 mediated stress response requires its translocation into nucleus which is dependent on actin[34, 35]. Nrf2 (Fig.4a) as well as vimentin (Vim) fluctuations (Fig.2 c) prompted an investigation into actin and other cytoskeletal elements. Upon closer inspection of lung proteome data *(Supplementary dataset S2)* we observed a striking effect on cytoskeletal elements due to altitude variation (Fig.5a). Across cytoskeletal proteins, we observed that 15K group had drastically lower cytoskeletal proteins while 25K (A) group showed recovery with protein levels being close to normoxic levels (BL) (Fig.5b). Moreover, the trinity of housekeeping proteins-actin (5,000 AU vs 15,000 AU), tubulin (17,000 AU vs 33,000 AU) (Fig.5b) and GAPDH (4,000 AU vs 7,000 AU) (Fig.6 c) were following the same trend. Immunoblotting confirmed the immense cytoskeletal and housekeeping perturbations in 15K group which are restored to normal in 25K (A) group. Since we had also observed declining serum albumin levels (0.294) (which bind Ca^2+^ ions) in 15K group lung *(Supplementary dataset S2),* we assayed Ca^2+^ levels in lung (Fig.5 c). As expected, free Ca^2+^ levels were the highest in 15K group (□5 mg/dL) while all other groups showed similar levels (1.5–2 mg/dL). This maybe one event when finely calibrated is causing a shift from 100% mortality to 100% survivability as its involved in various facets of cell signaling, intracellular trafficking and bio-energetics.

**Figure 5.**
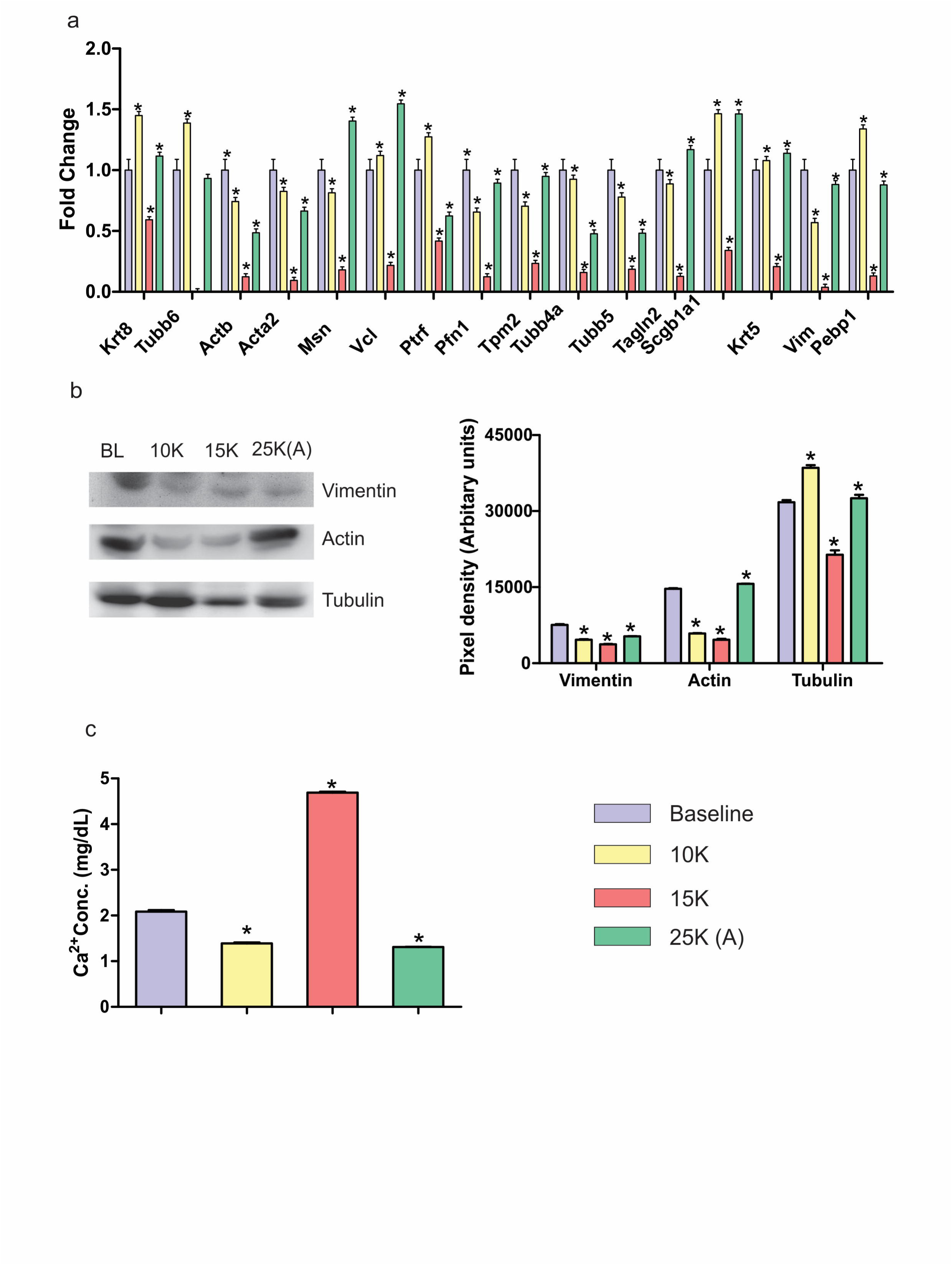
Cytoskeletal stability in 25K (A) group. a. Fold change values of various lung cytoskeletal proteins as observed in LC-MS/MS dataset. Across all cytoskeletal proteins, 15K group shows notable decline while 25K (A) group shows appreciable rebound.
b. Representative immunoblots of Vimentin, Actin and Tubulin in lung tissue. Vimentin levels decrease till 15K group with rebound in 25K (A) group. Actin levels decrease significantly in 10K and 15K groups with 25K (A) showing levels close to baseline. Tubulin levels also decline in 15K with rebound in 25K (A) group at a level close to baseline. Immunoblotting results are depicted as Mean±SEM of autoradiograms’ pixel intensities (Arbitrary units). Mean was calculated from three separate experimental replicates. * represents p-value<0.0001.
c. Bar graph representing free Ca^2+^ concentration in lung tissue across four groups. 15K group shows significant increase in Ca^2+^ concentration (4.7 mg/dL) as compared to baseline (2 mg/dL) while 10K and 15K groups (1.5 mg/dL approx) show slight decreases. Results are represented as Mean±SEM. Mean was calculated from three experimental replicates. * represents p-value<0.0001.

**Figure 6.**
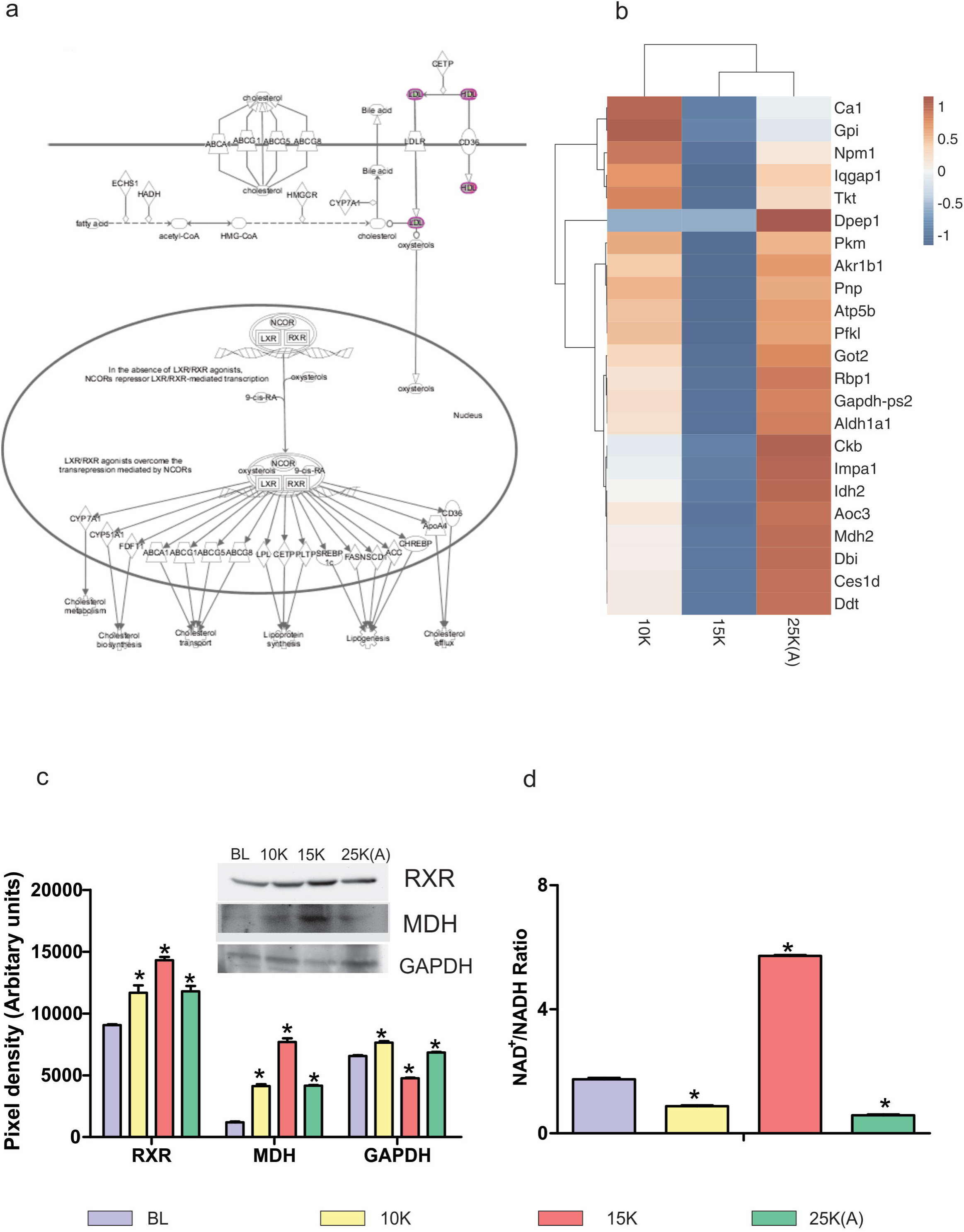
Restored energy homeostasis in 25K (A) group. a. Expanded view of LXR/RXR Activation pathway from IPA. The various downstream metabolic processes modulated by LXR/RXR such as cholesterol metabolism and lipogenesis as well as upstream molecules like Bile acid and oxysterols are shown.
b. Clustergram of 10K, 15K and 25K (A) with baseline fold change values taken as 1 for lung proteins involved in energy homeostasis and metabolism. Color brown indicates up-regulation while blue indicates down-regulation with white indicating neutral fold change. Fold change values were subtracted from baseline fold change (1) for each protein across all groups to arrive at final fold change values for clustergram. 15K and 25K (A) groups show complete anti-trends with 10K showing similarity with both the groups.
c. Representative immunoblots and respective bar graphs of Retinoid X receptor (RXR) and Malate dehydrogenase (MDH) in plasma and Glyceraldehyde 3-phosphate Dehydrogenase (GAPDH) in lung tissue. RXR and MDH (plasma) have maximal levels in 15K group with decline in 25K (A) group at levels similar to 10K group. GAPDH (lung) shows minimal levels in 15K group with increase in 25K (A) group to level close to 10K group. Immunoblotting results are depicted as Mean±SEM of autoradiograms’ pixel intensities (Arbitrary units). Mean was calculated from three separate experimental replicates. * represents p-value<0.0001.
d. NAD/NADH ratio was estimated in lung tissue. 15K group has highest ratio of NAD/NADH suggesting shift in pyruvate-lactate step towards lactate. 10K and 25K (A) groups show slight decrease in NAD/NADH ratio as compared baseline. Results are depicted as Mean±SEM. Mean was calculated from three experimental replicates. * represents p-value<0.005.

In the narrowest sense, calcium signaling effects in lung could be gauged in the plasma proteome by Vitamin D-binding protein’s (VDBP) trend and levels at varying altitudes. Recently, the Vitamin-D axis has been highlighted for lung diseases with emphasis on VDBP[36]. VDBP decreases significantly in 15K group (0.61) with recovery in 25K (A) group (0.95) *(Supplementary dataset S3).* VDBP/VDR axis highlights the bio-energetics aspect as VDR interacts with RXR[37].

### Glycolysis, TCA cycle and Lipid metabolism are affected in lung tissue: Maximal dysregulation in 15K group and recovery in 25K (A) group

LXR/RXR pathway (Fig.6a), due to its up- and down-regulation in plasma of 10K and 15K groups, respectively before normalizing again in 25K (A) group (Fig.2d) was investigated in context of bio-energetics. Also, lung proteome data *(Supplementary dataset S2)* revealed that most of the enzymes with roles in energy metabolism were unanimously down-regulated in 15K group while 25K (A) group experienced up-regulation (Fig.6b). RXR γ levels were analyzed to better understand not only calcium signaling related processes but the entire gamut of processes (Fig.6a) modulated by RXRs including lipid transport and reverse cholesterol transport[37]. We observed maximum RXR levels (□ 15,000 AU) in 15K group plasma (Fig.6 c) indicating an increased systemic stimulus for transcriptional activation while 10K and 25K (A) had similar levels (□ 12,000 AU). GAPDH, an essential housekeeping protein required for glycolysis, was found to decrease in 15K group lungs (□4,000 AU) (Fig.6 c). Malate dehydrogenase-1 (MDH-1) level which showed decline in 15K group lungs (0.345 fold change) *(Supplementary dataset S2)* was found to increase in 15K plasma (□8,000 AU) (Fig.6 c). This again lends credence to the hypothesis that systemic processes were activated when lung specific bio-energetic processes were dys-regulated. Finally to have an overview of the direction in which bio-energetics, particularly the pyruvate-lactate step was progressing, we assessed NAD^+^/NADH levels in lung (Fig.6d). It showed immense increase in 15K group (D6 AU) and a similar decline in 25K (A) group (C1 AU). This indicates a tendency for increased lactate production so as to regenerate NAD^+^ in 15K group lungs.

## DISCUSSION

Great efforts have been made to alleviate high-altitude maladies in terms of strategies like climb high and sleep low; supplemental oxygen; gamow bags and the plethora of medicines (nifedipine, sildenafil) specifically to reduce the patho-physiological effects of hypobaric hypoxia[38–43]. But the stunted growth of acclimatization protocols has fallen woefully short in providing a method for rapid induction/acclimatization to altitude. The current acclimatization schedule, stretching over weeks is not just time-intensive but also involves a greater economic cost and willpower to sustain. In this article, a proof of concept is posited that a shorter acclimatization strategy is not only feasible but can provide effective acclimatization against altitudes close to “death zone”. Secondly, protein networks and proteins that regulate/modulate the hypoxic response and provide a proteome based assessment of the acclimatization status of an individual are elucidated. This can be, after requisite testing and quality control in humans, the brother-in-arms for Lake Louise criteria.

Rapid induction to 25,000 ft represents a mortal threat to the organism. In our experiments and in real-world scenarios, one sees rat, mountaineers and aviators (pre-pressurized cabin era) struggle to circumvent mortality upon rapid induction[8]. In case of SD rats and mice, one sees a graded ascent to ≥25,000 ft (simulated) with multiple pre-exposures to lower altitudes for increased survival with a restricted ascent rate not beyond 200 m/min[44–46]. The nil survivability of 25K (D) group was worrisome and its circumvention but within minimum possible time form the core of the entire series of experiments. 25K (A) group was put through the shortest possible pre-exposure regimen (goal being rapid acclimatization) but the result was complete survival. In order to understand this complete reversal of mortality, we delved into lung tissue and plasma. For translational validity, we went for a highly trusted but low sensitivity; low throughput technique like MALDI-TOF[47] for plasma. The aim was to find the maximum number of perturbations in lung (LC-MS/MS) then understand it in terms of pathway analysis. Finally, the common links with plasma proteome signifying the effects of such exposures was to be determined with limited number of proteins. We observed 15K group protein levels were very low throughout in lung. In plasma, there was again a contradiction of sorts in 15K group and except some proteins (GPx3, STAT-3, RXR γ and MDH-1), all proteins either decrease or remain at similar levels as 25K (A). When we further investigated the various oxidative stress and antioxidant parameters biochemically, we observed that maximum damage is occurring in the lung and maximum systemic activation of redox homeostasis mechanisms is there in plasma. Thus, only upon failure of lung redox homeostasis is there activation of systemic redox homeostasis. Very high altitude zone causes an intense redox homeostasis upheaval. But as observed through pathway analysis, there are not many overall up- or down-regulations in canonical pathways. In the real world also, there is no mortality observed in 15K group. Thus, we can safely state that there is a major systemic activation in 25K (A) group of systemic redox homeostasis mechanisms. Oxidative stress specific lung transcripts also indicate a heightened oxidative stress response in 15K group while 25K (A) group responds in a manner strikingly similar to 10K and normoxic groups. This is due to pre-exposure at 15,000 ft (very high altitude) for 10 h followed by normobaric normoxia exposure for 1 h. Central to redox homeostasis modulation are the glutathione and thioredoxin reserves. One observes that respective peroxidases and reductases related to these two also play very important and crucial roles in countering the oxidative stress and maintaining redox homeostasis. An interesting feature is the rise of both GSH and TRX reductase levels in 25K (A) group lung tissue suggesting recovery of redox balance while in plasma there is a concomitant decrease in levels of both in 25K (A) group. This anti-trend between same/related molecules in lung and plasma strongly indicates that hypobaric hypoxia elicits a systemic redox homeostatic response upon the overwhelming of lung redox homeostasis. Such anti-trends bestow us with a calibration with which to diagnose the acclimatization status of an organism by simply analyzing their plasma. This aspect is further elucidated in terms of a network between lung and plasma and will be elaborated in later paragraphs (Fig.7).

**Figure 7.**
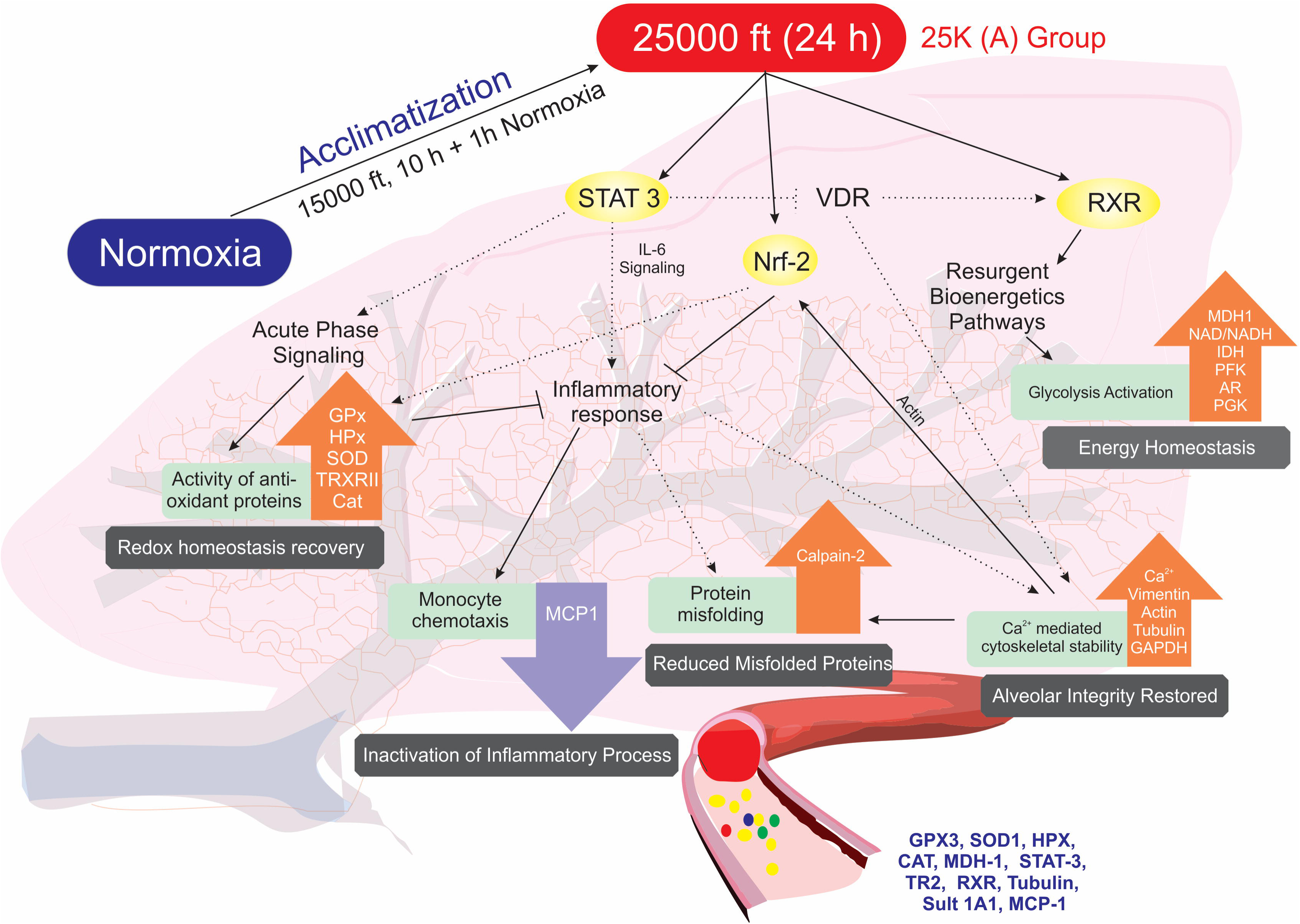
Putative protein cascades involved in rapid acclimatization in 25K (A) group. Figure represents protein networks and processes involved in redox homeostasis, energy homeostasis, inflammatory signaling, protein misfolding and cytoskeletal (alveolar) integrity for 25K (A). Dotted black lines represent events derived from literature. Bold black lines represent experimental findings. Central signaling molecules like STAT-3, Nrf and RXR begin cascade. STAT-3 levels fall back to normoxic levels in 25K (A) group plasma causing lowered inflammatory signaling, increased VDR/RXR signaling and increased anti-oxidant response. Lowered inflammatory signaling is observed via MCP-1 levels as well as increased Calpain-2 levels. Increased VDR/RXR signaling helps restore energy homeostasis in 25K (A) group as compared to 15K group which is observed via NAD/NADH ratio and increased levels of glycolytic enzymes. VDR also plays a small role in calcium homeostasis. Free Ca^2+^ levels are reduced significantly in 25K (A) group. This helps restore levels of cytoskeletal proteins with housekeeping functions. This, in part, causes alveolar structural integrity to endure extreme hypobaric conditions. Reduction in free Ca^2+^ levels also improves Calpain-2 levels. Actin is required for Nrf2 translocation into nucleus. When levels of actin are restored, Nrf2 mediated oxidative stress response is activated. Inflammatory signaling is further reduced due to it. Also, antioxidant protein levels increase to strengthen redox homeostasis. This entire process occurs within lung and plasma. Thus, proteins like GPX3, SOD1, HPX, CAT, MDH-1, STAT-3, TR2, RXR, Tubulin, Sult 1A1 and MCP-1 can provide important clues regarding the acclimatization status when their levels at normoxia and extreme hypobaric hypoxia (as in 25K (A) group) are compared on following the stated acclimatization strategy of 10 h at 15,000 ft followed by 1 h of normobaric normoxia exposure.

We observed acute phase signaling to play a major role in the initial response to hypobaric hypoxia alongwith redox homeostasis. Shared molecules in both these pathways like hemopexin play a major role in subverting any major damage during the exposure. An interesting feature of this entire study is the involvement of calcium signaling that is expressed in plasma as VDBP fluctuations. Such fluctuations as well as lung tissue specific calcium signaling dependent molecules like Annexins coupled with changes in Ca2+ levels indicate a thorough involvement of calcium signaling in actuation of hypoxia acclimatization. Since the acute phase signaling also consists of molecules involved in inflammatory processes, we measured not just the levels of proteins observed to fluctuate such as C3 (Supplementary) but also of related effectors like MCP-1. The results were clearly indicative that at very high altitudes, we have a severely pro-inflammatory milieu but in 25K (A) group in spite of the extreme altitude we see a huge downfall in MCP-1 levels. This is in complete contradiction to previous studies where hypobaric hypoxia exposure triggers inflammation[48–50]. The pre-exposure regimen is the only plausible reason to explain the dramatic decrease in pro-inflammatory signals at a much higher intensity of hypobaric hypoxia. Hence, we can safely commit that the animals are saved from any patho-physiologies as a function of downregulated inflammatory processes.

Altitude variation also affects essential cytoskeletal elements and housekeeping proteins. Actin, tubulin and GAPDH levels were found to be drastically reduced in 15K group lungs. They rebound to close to normoxic levels in 25K (A) group but a fluctuation is observed across the four groups. This is an event we suppose could be causing the lung tissue to fail at extreme altitude as cytoskeletal re-arrangements will affect every single aspect of lung cell functions from metabolism to signaling to endocytosis. This particular event has been noted to be prominent in HAPE cases. HAPE patients also suffer from major re-shuffling and disruption of cytoskeletal elements, particularly actin[51] evident via increased lung permeability[52–54]. The restoration of these elements close to normoxic levels in 25K (A) group is again an assurance of the robustness of our pre-exposure regime. Hence, the rebound of all cytoskeletal elements observed in 25K (A) group at levels similar to normoxic controls indicate a termination of events that may lead to HAPE at least at the cellular level.

HAPE patients whose metabolome was analyzed were found to have dysregulated glycolysis and upregulated protein catabolic processes [55, 56]. The analysis of lung proteome also revealed extensive perturbations in metabolic pathways and bio-energetics. We observed multiple enzymes of the glycolysis, TCA and lipid metabolic networks to be severely reduced in 15K group indicating a dysregulated metabolism. Also, ATP synthase sub-units were found to be severely marred in the same group indicating inefficient bio-energetics. Upon the pre-exposure, the 25K (A) group shows recovery on all these counts which are carried over into the plasma proteome in spite of increased intensity of hypobaric hypoxia. The restoration of the bio-energetics process in 25K (A) group is another ace indicator of the robustness of this process of rapid induction to altitude.

In terms of a network, as visualized in the concluding figure (Fig.7), we observe STAT-3, RXR γ and Nrf-2 as messengers modulating the response of various downstream proteins involved in the acute phase response, inflammatory signaling, metabolic processes and cytoskeletal stability. STAT-3 levels decline in 25K (A) group to allow for recovery of RXR which stabilizes downstream processes leading to energy homeostasis. Declining STAT-3 also helps reduce inflammatory stimuli via IL-6 signaling. Although there was no significant change in C3 levels across groups we observe MCP-1 (chemoattractant for monocytes) levels to fall back to normoxic levels after dramatic rise in 15K group. Also, increasing Calpain-2 subunits reduce levels of redox stress induced misfolded peptides and proteins. As a retrospective effect, we also observe declining free Ca^2+^ levels in lung and restoration of certain essential cytoskeletal elements like actin, tubulin and vimentin. Thus, cytoskeletal stability is restored in 25K (A) group. The restoration of energy homeostasis and cytoskeletal stability is testament to the effectiveness of the acclimatization regimen. However, without the restoration of redox homeostasis both these processes are but temporary relief measures against hypoxia. In this regard, we observe Nrf-2 mediated oxidative stress response as well as STAT-3 decline cause a resurgence of anti-oxidant proteins in the lung. The signature of these processes in the lung can be assessed in the plasma via the protein levels of the mentioned proteins (Fig.7) in plasma. The anti-trends revealed earlier in redox homeostasis (biochemical assays, immunoblots) in lungs and plasma must be accounted for while assessing the panel of plasma proteins shown. In order to make sense out of the various proteins in the panel, all of them must be assessed at baseline, very high altitude and extreme altitude zones. This shall provide a comparative analysis of protein trends. Similar trends as described in this article will indicate a positive acclimatization status while a negative acclimatization status will be described as opposing protein trends. Proteins with potential to detect acclimatization status of individuals going through the stated acclimatization strategy are glutathione peroxidase 3 (GPX3), superoxide dismutase 1 (SOD), hemopexin (HPX), catalase, malate dehydrogenase 1 (MDH-1), STAT-3, thioredoxin reductase 2 (TR2), retinoid X receptor (RXR), tubulin, sulfotransferase 1A1 (Sult 1A1) and monocyte chemoattractant protein 1 (MCP-1) (Fig.7).

### Study Limitations

Samples from the 25K (D) group were unavailable for comparative analysis with other groups due to widespread mortality in this group. The molecular occurrences that signify the progression of patho-physiological processes ultimately leading to death due to hypobaric hypoxia remain occluded. Also, we have not forayed into chronic exposures after this acclimatization regimen. This we feel is a subject for future research and publications. Trials using human subjects for rapid acclimatization will require further scientific diligence. This can also be a fruitful future endeavor. In spite of the authors’ best efforts, these limitations do remain.

### Conclusion

To conclude, our data proves that exposure to very high altitude (15,000 ft for 10 h) followed by a short normobaric normoxia exposure (1 h) causes a 100% shift from mortality to survival during further acute exposure (24 h) to extreme altitude of 25,000 ft. This study suggests the possibility of a new route of rapid acclimatization, which was found to bring a high degree of survival at extreme altitude (25,000 ft) by pre-exposure to 15,000 ft, which otherwise showed zero survival rate in simulated condition. We had previously investigated and found a high degree of similarity between rat and human regarding pathways and networks responding to hypobaric hypoxia[57]. This study may therefore stand as a proof of concept to translate the objective of rapid acclimatization without compromising the safety and performance of an organism during extreme hypobaric hypoxia exposure. Secondly, this study also provides a putative panel of plasma proteins to objectively state whether the organism is acclimatizing rapidly to extreme altitude upon using the stated acclimatization strategy. Rapid acclimatization, as a process, can potentially thwart many hazards and disasters (natural and man-made) in high-altitude regions. When translated, rapid acclimatization will save many medical contingencies.

## Materials and Methods

### Experimental Animals & Experiment Design

Age matched (10 week old) male Sprague Dawley rats weighing 200–230 g were used for all the experiments. The animals were housed in polypropylene cages with paddy husk as substratum at 25±5˚C with humidity at 50±5%. Night-day cycles of 12 h each was maintained. Each cage held 3 rats with provisions for food and water. Animal procedures and experimental protocols were approved by Institutional Animal Ethics Committee (Authorization Number: 27/1999/CPCSEA) and we followed the standards set forth in the Guide for the Care and Use of Laboratory Animals (National Academy of Science, Washington, D.C.).

Forty-five rats (n=45) were randomly and equally divided into five groups, first Baseline controls (animals not receiving simulated hypobaric hypoxia exposure), 10K (animals exposed to simulated hypobaric hypoxia equivalent to 10,000 ft/ 521 mmHg for 24 h), 15K (animals exposed to simulated hypobaric hypoxia of 15,000 ft; 429 mmHg for 24 h), 25K (A) (animals receiving a pre-exposure at 15,000 ft for 10 h followed by normobaric normoxia exposure of 1 h prior to exposure of 25,000 ft; 282 mmHg for 24 h, A stands for Acclimatized) and 25K (D) (animals receiving simulated hypobaric hypoxia exposure of 25,000 ft; 282 mmHg for 24 h, D stands for direct exposure, without acclimatization). There was no pre-acclimatization protocol followed for any groups except in case of 25K (A). Triplicates were run for each of the five groups with three animals in each run per group.

### Hypobaric Hypoxia Exposure and sample collection

Hypobaric Hypoxia exposure simulation was performed in custom designed hypobaric hypoxia simulation chamber (7 star systems, Delhi, India). The chamber had a constant temperature and humidity of 25±5˚ C and 50±5%, respectively and an ascent rate of 587 m/min was maintained during the exposure. Airflow of 2 l/min was also maintained in the chamber. Animals were provided with food and water inside the chamber. Immediately after completion of exposure, all animals were sacrificed using approved procedures. The blood was collected in EDTA containing tubes which were further processed to obtain plasma. Protease inhibitor cocktail (Cat # P8340, Sigma, USA) was added to prevent protease activity. Lung samples were rinsed in cold PBS and snap frozen in liquid nitrogen. All the samples were stored in -80°C until further use.

### Biochemical assessment of oxidative stress and antioxidant levels

#### ROS estimation using DCFHDA

A non-fluorescent lipophilic dye, Dichlorofluorescein diacetate (DCFHDA) (Cat # C400, Life Technologies, USA), was used to measure ROS levels in both lung tissue homogenate and plasma of all four groups. Once internalized into the cells, it is cleaved into 2,7-dichlorofluorescein by intracellular esterases and on combining with ROS cleaved DCF produces fluorescence. The fluorescence produced is directly proportional to the ROS levels. We added 10 µl of 10 μM DCFHDA to 150 μl of lung tissue homogenate (10% w/v in RIPA buffer, Thermo Fisher Scientific, Waltham, USA) and incubated for 40 minutes at 37°C in amber tubes (Borosil, USA) in the dark. For estimation of ROS levels in plasma, samples were diluted 1:30 in RIPA buffer and then 150 µl of this diluted plasma was incubated with 10 µl of 10 μM DCFDA at 37˚C for 40 minutes in dark amber tubes. Finally, fluorescence was measured at 488 nm excitation and 525 nm emission wavelengths, respectively using flourimeter (LS45 Luminescence Spectrometer, PerkinElmer, USA). The florescence units were normalized to background and data was then presented as AU per µl of sample.

#### Lipid peroxidation status using MDA Assay

The method suggested by Ohkawa et al was used to measure lipid peroxidation status in both lung tissue homogenate and plasma [58]. Briefly, 750 μl of trichloroacetic acid (TCA; 20% w/v in distilled water) and 750 μl of thiobarbituric acid (0.67% w/v in 0.05 M NaOH) were added to 250 μl of the lung tissue homogenate and plasma each, in series, incubated in a water bath at 95°C for 15 minutes then allowed to cool to room temperature. The mixture was then centrifuged (400 *g*; 5 minutes). 200 μl of the supernatant from each group was added in triplicate in a 96-well plate and optical density was measured at 531 nm using spectrophotometer (VersaMax ELISA Microplate Reader, Molecular Devices, Sunnyvale, CA, USA). The data was normalized with protein content of the samples estimated using standard Bradford’s assay and subsequently represented as μmol MDA/mg of protein

#### Estimation of Superoxide Dismutase activity

Quantification of Superoxide dismutase activity was accomplished using Enzychrome^TM^ superoxide dismutase assay kit (Cat # ESOD-100, Bioassay systems, USA) as per the manufacturer’s instructions. In concise form, 20 µL standard or plasma/lung tissue homogenate samples were added in triplicate to a 96 well plate. Then 160 µL working reagent containing assay buffer, xanthine and WST-1 were added to each well in given order. Optical density was measured immediately at 430 nm (OD_0_) and then plate was incubated at 25°C for 60 minutes. Again, optical density was measured at 430 nm (OD_60_). Finally the concentration of SOD was measured using ΔΔOD vs SOD concentration standard curve. Activity was represented as U/ml.

#### Estimation of Reduced Glutathione

Reduced glutathione was measured using EnzyChrom^TM^ GSH/GSSG Assay kit (Cat. No. EGTT-100, BioAssay Systems, USA) in both lung and plasma samples. Briefly, 25 μl lung homogenates and plasma samples from each group were deproteinated using 65 μl of 5% wt meta-phosphoric acid. About 12 μl of clear supernatant is taken and mixed with 488 μl of 1X Assay buffer. Then, 200 μl of the prepared sample was added to a single well of ELISA 96-well plate and 100 μl Working Reagent (1X Assay buffer, GR enzyme, NADPH & DTNB) was subsequently added. Absorbance was taken immediately and after 10 min at 412 nm.

#### Thioredoxin reductase assay

Thioredoxin reductase was measured using TRX Reductase assay kit (Cat. # CS0170, Sigma Aldrich, USA) as per the manufacturer’s instructions. Briefly, 7 μl of either lung tissue homogenate or plasma samples were taken and 7 μl Assay buffer, 180 μl wash buffer and 6 μl dithio-bis-nitrobenzoic acid (DTNB) were added in sequence. 4 μl of dilute inhibitor solution was added to a duplicate set of samples, the rest of the protocol being identical for both sample sets. Absorbance was measured for the two duplicate sets every minute at 412 nm. The rate of change in absorbance per unit time (ΔA_412_/min) for the set with inhibitor solution was deducted from the sample set without inhibitor to obtain the final ΔA_412_/min. the change in absorbance was plotted as TRX reductase levels in the sample.

#### Total calcium estimation

Calcium concentration was estimated in both lung tissue homogenate and plasma using Quantichrom Calcium assay kit (Cat # DICA-500, BioAssay Systems, USA), as per the manufacturer’s instruction. Briefly, 5 μl of either plasma and lung samples or standard was added in duplicates to a 96-well plate. Then, 200 μl of working reagent was further added and the well contents were mixed by tapping. The plate was incubated for 3 minutes at room temperature, the absorbance was then immediately recorded at 612 nm.

#### NAD/NADH assay

NAD/NADH ratio was measured in lung tissue using EnzyChrom™ NAD/NADH Assay kit (Cat # E2ND-100, BioAssay Systems, USA) as per manufacturer’s instructions. Briefly, two tissue samples from same animal, weighing 20–25 mg each, were taken and homogenized in 100 µl of NAD or NADH extraction buffer, respectively. The extracts were heated for 5 min at 60 ˚C and 20 µl assay buffer followed by 100 µl of the opposite extraction buffer were added in sequence. The samples were vortexed and centrifuged (14000 rpm, 5 min) and clear supernatant was used for the assay. Each well had 40 µl sample or standard and 80 µl Working reagent. Absorbance was measured at zero and 15 min interval at 565 nm at room temperature. Ratio was calculated using the manufacturers’ equation.

### High throughput proteomics of lung tissue using LC-MS/MS

#### Sample preparation

Equal mass of lung tissues excised from similar regions from the representative animals of each experimental group (N, 10K, 15K, 25K) were homogenized on ice in Radioimmunoprecipitation assay (RIPA) buffer (5 ml). Homogenised tissues were then centrifuges at 15000 g for 10 min to separate precipitated debris and supernatant was collected. The pellet was further treated with Protein Isolation buffer (ToPI) and centrifuged. The lysate formed was added to the retained supernatant. Quantity of protein in each sample was estimated using Bradford assay. 100 µg of protein from each sample was dispensed and processed for MS. Processing involved reduction, alkylation and finally precipitation for removal of interfering substances. Then, trypsin digestion was performed overnight. The digested peptides in digestion buffer were labelled with iTRAQ reagents. Each sample was SCX fractionated and the fractions eluted at 75 mM, 150 mM, 450 mM ammonium acetate were collected and analyzed individually by nano-LC-MS/MS. The combined data was used for MudPIT.

#### Mass spectrometry (LC-MS/MS)

Desalting was done using ZipTip and then the samples were speedvac dried before re-suspension in mobile phase for LC-MS/MS. Peptides were eluted from the column using a linear acetonitrile gradient from 5 to 45% acetonitrile over 180 minutes followed by high and low organic washes for another 20 minutes into an LTQ XL mass spectrometer (Thermo Scientific) via a nanospray source with the spray voltage set to 1.8kV and the ion transfer capillary set at 180 ºC. A data-dependent Top 5 method was used where a full MS scan from m/z 400–1500 was followed by MS/MS scans on the five most abundant ions. iTRAQ Ratio > 1.5 are classified as up-regulated, < 0.67 are classified as downregulated. Ratios from 1.5 – 0.67 are considered moderate to no changes.

### Plasma proteomics using two dimensional gel electrophoresis (2DGE)

#### Protein separation by IEF and SDS-PAGE

Isoelectric focusing was performed with Immobiline Dry Strip, pH 4–7, 18 cm (GE Healthcare, Sweden) on IPGphor IEF System (GE Healthcare, Sweden) at constant voltage. The strip was pre-incubated with 350 µl rehydration buffer containing 7 M urea, 2 M thiourea, 1.2% w/v CHAPS, 0.4% w/v ABS-14, 20 mM dithiothreitol (DTT), 0.25% v/v pH 3–10 ampholytes, 0.005% w/v bromophenol blue (BPB) and 300 μg protein at room temperature for 18 h. The IEF consisted 500 V for 7 h (slow), 1000 V for 1 h (linear), 8000 V 3 h (gradient), 8000 V 3 h (linear), 10000 V 2 h (gradient) and 10000 V 1 h (linear). Prior to the second-dimensional gel separation, the IPG strips were equilibrated for 2 x 15 min with gentle shaking in 6 ml of SDS equilibration buffer [50 mM Tris–Cl (pH 8.8), 6 M urea, 30% v/v glycerol, 2% SDS]. Freshly prepared DTT (2%, w/v) was added in the first step and iodoacedamide (2.5%, w/v) in the second equilibration step. The second dimension was carried out using EttanDaltSix Electrophoresis System (GE Healthcare, Sweden). The strips were then loaded onto 12% SDS-polyacrylamide gel and sealed with 0.5% agarose (containing BPB). The running buffer contained 25 mM Tris–HCl, pH 8.3, 192 mM glycine and 0.1% w/v SDS. Electrophoresis was performed at a constant current of 25 mA per gel at 25°C for 6 h.

#### Silver staining of 2D gels

After gel electrophoresis, proteins were visualized by modified silver staining procedure compatible with MS. The gels were fixed in 50% v/v methanol, 12% v/v acetic acid and 0.05% v/v formaldehyde for 2 h. The fixed gels were then rinsed with 50% v/v ethanol three times for 20 min each, then again sensitized with 0.02% w/v sodium thiosulfate followed by three washings with Milli-Q water each for 20 seconds. The gels were immersed in 0.1% w/v silver nitrate and 0.075% v/v formaldehyde for 20 min and rinsed with Milli-Q water twice for 20 second each followed by developing with 6% sodium carbonate and 0.05% v/v formaldehyde. Finally, the reaction was terminated by adding 12% v/v acetic acid.

#### Image acquisition and data analysis

The stained gel images were captured using an Investigator™ProPic II (Genomics Solutions, UK) and the digitized gel images analyzed using 2D-Progenesis Samespot Software (Non Linear Dynamics, USA). Spot matching between all the gels were viewed using the automatic spot detection and normalization tool and edited where appropriate. The relative intensity of individual spots in 2D-GE gels of different time exposure, treatment and control rats were quantified using a gray-scale and the differences between spot pairs were determined. The median differences of relative spot intensities in matched spots were calculated. Protein alterations confirmed in at least four sample pairs were scored as significant.

#### In gel tryptic digestion of proteins

In gel tryptic digestion of the proteins was performed using 20 ng/µl sequencing grade trypsin, (Cat no.V5111, Promega, USA) protein digestion, silver stained 2-D gels were extensively washed twice at for 10 min each with MilliQ water and differential spots were manually excised and subjected to in-gel digestion. The gel pieces were de-stained in a freshly prepared 1:1 solution of 30 mM potassium ferricyanide and 100 mM sodium thiosulfate for 1–2 min until the brownish colour disappeared. The gel pieces were then quickly rinsed thrice with MilliQ water to stop the reaction. Next, the gel pieces were further washed with 50 mM ammonium bicarbonate/acetonitrile for 15 min at room temperature. Enough acetonitrile were added to cover gel pieces for shrinking the gel pieces. The gel pieces were rehydrated with 10 mM ammonium bicarbonate for 5 min, followed by incubation with equal volume of acetonitrile for 10 minutes. The gel pieces were again covered with acetonitrile to induce shrinkage and subjected to vacuum drying. The dried gel pieces were treated with 20 µl of sequencing grade trypsin overnight at 37°C. The tryptic digested peptides were sonicated for 10 min and dried in MAXI dry plus (Heto Holton, UK).

#### Matrix preparation

For identification of proteins, CHCA matrix mixed in 50% ACN in 0.1% TFA. After mixing, solution was sonicated for 10 minutes and centrifuged at 10000 rpm for 2 minutes. Clear supernatant was used as saturated matrix for spotting.

#### Protein Identification by MALDI-TOF/TOF

For PMF, in-gel tryptic peptides were mixed with CHCA matrix and exposed to laser radiation. The matrix was prepared using 70% acetonitrile and 0.03% TFA. 0.5 ml each of peptide extract and matrix were mixed and manually spotted onto 384 well AnchorChip sample target (Bruker Daltonics) and dried at room temperature with surface cover. Peptide mass spectra were recorded using an Ultraflex III Tof/Tof mass spectrometer (Bruker Daltonics) with a 384-sample scout source in reflectron mode. The ion acceleration voltage after pulsed extraction was 27000 V. All data was recorded automatically on the MALDI-TOF/TOF instrument using the three most abundant peptide signals of the corresponding peptide mass fingerprint (PMF) spectrum. The monoisotopic peak list was generated in Post Processing s/w and True peptide mass list was generated by Bruker Flex Analysis software version 3.0 and Biotools ver 3.1 without using the smoothing function and the signal to noise ratio of 20 was set. The generated peptide mass list was searched with MASCOT (http://www.matrixscience.com) using entire Uniprot/Swiss-Prot protein database to find and match the identified protein. Databases searches were performed by using following search parameters; Rattus norvegicus as taxonomy, carbamidomethyl modification of cysteines and possible oxidation of methionine, one missed cleavage, a mass accuracy of #100 ppm was requested for PMF and for MS/MS searches. For each identified Protein, at least one Peptide was selected for MS/MS (TOF/TOF) to validate the Protein Identity. Instrument was used in Lift mode (TOF/TOF) to obtain MS/MS spectra. Again the Flex Analysis3.0 and Biotools 3.1 s/w were used to generate the fragments mass list and the sequence Tag of peptide. The mass list was sent to database in same way as was done in case of above PMF approach. The mass tolerance error of 0.5 Da to 1.0 Da was used for MS/MS ion search. The MS/MS ion search confirmed the protein identity and provided the amino acid sequence of particular peptide.

### Network analysis of Proteomics data

Data obtained from both high throughput LC-MS/MS and 2DGE-MS was curated manually for artefacts and sorted according to the false discovery rate (less than 1%) and p value (less than 5%). Selected lists of proteins with respective fold change were imported into Network analysis tool Ingenuity Pathway analysis (IPA, Qiagen) with inbuilt statistical analysis package. With the set fold change cutoff of 1.3 various integrated programs such as canonical pathways, network of diseases and functions, proteins networks were used to determine key changes in the cellular events, biochemical processes and molecular cascades. A significant positive and negative z-score was used to predict the directionality of the cellular event. Top canonical pathways and molecular events were selected on the basis of minimum p-value.

### ELISA

ELISA assays were performed for MCP-1 (make, model), regucalcin (make, model) and Sulfotransferase 1A1 (make, model). In case of MCP-1 and Regucalcin, the assays were performed in strict adherence of the manufacturer’s protocol for plasma and lung tissue homogenate, respectively while in case of Sulfotransferase 1A1, both plasma and lung samples were assayed separately as per the manufacturer’s guidelines.

### Immunoblotting

Immunoblotting of proteins were performed in the lung tissue homogenate samples and plasma samples. Sample containing 25 µg of total protein were separated on 10% SDS-PAGE gels and transferred onto nitrocellulose/PVDF membrane. The membrane was blocked in 5% skim milk in PBS-0.1% Tween-20 (PBST) overnight at 4 °C. Further, the membrane was washed three times with PBST for 10 min each and incubated with their respective primary and secondary antibodies for 2 h at room temperature. After washing, the membrane was developed by adding chemiluminiscent peroxidase substrate (Cat # CPS1300, Sigma, USA). Densitometric analysis of autoradiograms was performed using Image J software (http://rsbweb.nih.gov/ij/).

### PCR Array

#### RNA isolation and cDNA synthesis

Transcription of antioxidant specific genes was evaluated with quantitative real-time qRT-polymerase chain reaction (PCR) in four experimental groups-Normoxia, 10K, 15K and 25K. Total RNA was isolated from lung tissue using TriReagent as per manufacturers’ protocol. Quantification of RNA was done using Biophotometer plus, Eppendorf and RNase-free water. Reverse transcription was initiated promptly after quantification of RNA using High capacity c-DNA Reverse Transcription kit (cat no.4368814, ABI) as per the manufacturer’s instructions. The synthesis program appended a single incubation at 42 °C for 15 min, followed by incubation at 95 °C for 5 min. The reaction volume was made up to 100 μl with MilliQ water. This cDNA was used for Quantitative RT-PCR.

#### Real time PCR

Rat RT2 Profiler ™ PCRArray profiling the expression of 84 genes related to oxidative stress, ROS metabolism and related oxygen transporter genes from Qiagen (PARN 065Z, Qiagen, USA) was used for analysis of oxidative stress specific transcripts as per the manufacturer’s guidelines. Briefly, 1173 µl nuclease-free water (AM9937, Ambion, USA), 1150 µl SYBR green (ABI) and 100 µl cDNA were mixed in a solution. 25 µl was added to each well of the 96-well array plate from this solution. This plate was briefly spun for 20 sec at 1100 rpm. RT-PCR protocol was set at 95 ˚C for 10 mins. Then 40 cycles were performed with 95˚ C for 17 seconds and 60˚ C for 1 min. Applied Biosystems StepOnePlus was used for performing the RT-PCR.

#### Data Analysis

Analysis was done based on the ΔΔC_T_ values. Fold change values for each gene were tabulated and further analysed using Qiagen’s RT-PCR data analysis tool (http://pcrdataanalysis.sabiosciences.com/pcr/arrayanalysis.php). Tool provided a quality check step followed by normalization of data with housekeeping genes and then determination of statistically significant up and down-regulated genes. The data was plotted as dotplot and heatmap of individual comparison, additionally a heatmap with hierarchical clustering was also obtained using the manufacturers’ online support tool. Additionally, the data was finally analysed using Ingenuity pathway analysis as stated in the protein analysis.

### Statistical analysis

Data sets were analyzed using Graph pad prism (v 5.0). For statistical significance, 1-way analysis of variance (ANOVA) followed by Bonferroni’s Multiple Comparison test was applied. Significance was set at p<0.05. In all cases, three independent experiments were carried out. Results were presented as mean value ± standard error of mean (SEM).

## Acknowledgements

This study was funded by the Defence R&D Organization, Ministry of Defence, Govt of India under the project DIP-263. Subhojit Paul is a recipient of Senior Research Fellowship, CSIR. Anamika Gangwar is a recipient of Senior Research Fellowship, DST-INSPIRE. Authors acknowledge Dr. Aditya Arya for his creative inputs in designing the concluding figure. Authors also acknowledge Dr. Shantanu Sengupta and Swati Varshney for access to MALDI-TOF facility.

## Author Contributions

Experiments were conceived by YA and SP. Experiments were performed by SP and AG. Representative figures were designed by SP and AG. Manuscript was written by YA and SP. KB critically evaluated the manuscript.

